# Divergence in skeletal muscle growth by differential spatial hyperplastic patterning in teleost fishes

**DOI:** 10.64898/2026.02.23.707519

**Authors:** Yansong Lu, Marco Podobnik, Koji Ando, Michael Pan, Jake Locop, Angus Guo, Philippe Mourrain, Kazu Kikuchi, Avnika A. Ruparelia, Peter D. Currie

## Abstract

Skeletal muscle plays important locomotive and metabolic functions, yet its formation and maintenance are processes remaining largely unclear mechanistically in any animal. Teleost fishes display extraordinary muscle growth due to their ability to undergo both hyperplasia and hypertrophy throughout life. These phenomena vary greatly even between closely related species, providing opportunities to elucidate growth dynamics and underlying mechanisms through cross-species comparisons. Using histological and genetic approaches, we compared muscle growth dynamics in three closely related danionin species with distinct growth capacities: the giant danio (*Devario malabaricus*), the zebrafish (*Danio rerio*), and *Danionella cerebrum*, as well as the more distantly related African turquoise killifish (*Nothobranchius furzeri*). Our study reveals alterations in spatial patterning of muscle hyperplasia and developmental timing to be major contributors to observed differences in muscle growth between examined species. Single-cell RNA profiling, in situ hybridization chain reaction and cell type-specific mutagenesis revealed muscle stem cell-specific expression of extracellular matrix genes that mediate stem cell activity, which in turn may drive growth differences between species. Taken together, our findings highlight autonomous regulation of muscle stem cells as a conserved but adaptable mechanism governing muscle patterning and diversification.

## Introduction

In vertebrates, skeletal muscle plays indispensable roles in supporting locomotion and metabolism. In humans, loss of muscle mass due to genetic conditions or aging not only drastically reduce quality of life, but is also an unfavorable prognostic factor for many disease and conditions, including cancer^1–5^, infections^6–9^, organ transplant^5,10,11^, as well as overall longevity^12–14^. Despite its critical importance, muscle mass in humans is difficult to maintain, especially with increased age^15,16^. Teleost fishes have the remarkable ability to maintain or even increase their muscle mass with age^17^. The genetic mechanisms that drive teleost muscle stem cells (MuSCs) to form myofibers lifelong remains to be fully elucidated, but understanding these mechanisms could provide a new avenue for enhancing muscle mass in human patients and lead to novel therapeutic intervention to combat muscle aging in humans.

Skeletal muscle growth is primarily facilitated by two distinct processes – hyperplasia and hypertrophy. While hyperplasia involves the formation of multinucleated myofibers through differentiation and fusion of MuSCs – also known as myoblast fusion^18^, hypertrophy defines the enlargement of existing myofibers through protein synthesis, as well as the fusion of additional MuSCs into existing myofibers to meet the increased transcriptional demand of the enlarged myofibers^19^. Due to the essential role of MuSCs in supporting both modes of muscle growth, they are regulated by elaborate mechanisms to ensure precise control of skeletal muscle growth^20–22^. These processes can be manipulated for therapeutic gains, though the underlying regulatory mechanisms remain to be fully elucidated.

Modes of hyperplasia and hypertrophy differ drastically between amniotes and teleosts – while mammalian hyperplasia is restricted to prenatal stages, with postnatal muscle growth being solely by hypertrophy, teleost skeletal muscle undergoes both hyperplasia and hypertrophy throughout life, which enables extraordinary growth capacity in teleosts^23,24^. Hyperplastic muscle growth capacities vary drastically even between teleost species, with some species portraying significant lifelong hyperplasia, whereas others experience sharp growth plateaus similar to mammals^25–29^. This divergence creates an opportunity to understand the dynamics and mechanisms underlying hyperplastic muscle growth in teleosts through cross-species comparison.

Here, we performed histological and transcriptomic characterization of skeletal muscle growth and MuSC dynamics in three closely related danionin species that portray drastically different growth capacities, including the giant danio (*Devario malabaricus*), the zebrafish (*Danio rerio*) and *Danionella* (*Danionella cerebrum*, hereafter: *Danionella*). Additionally, we examined the African turquoise killifish (*Nothobranchius furzeri*), which is evolutionarily distant to the danionins yet shares many similarities in muscle growth dynamics to certain danionin species. By leveraging a newly developed tool for high-throughput morphometric analysis, we revealed divergent hyperplastic muscle growth capacities between species, including drastic alterations in spatial patterning and developmental timing (heterochrony). For example, while muscle hyperplasia in zebrafish and giant danio occurs evenly across the myotomes, hyperplasia deep inside the myotome is greatly reduced in killifish and completely absent in *Danionella*. Using single-cell RNA sequencing (scRNA-seq) and in situ hybridization chain reaction (HCR), we revealed important roles of MuSC-expressed extracellular matrix (ECM) genes in regulating stem cell activation and differentiation across species. Additionally, increased MuSC abundance was observed by MuSC-specific deletion mutations in a collagen type IV gene in juvenile zebrafish. Our results suggest that these stem cells regulate their activity autonomously by altering ECM composition and that divergence in MuSC-specific ECM gene regulation may contribute to the different growth capacities observed between species. Our study sheds light on the regulation of hyperplasia as a conserved but adaptable mechanism driving skeletal muscle growth in teleost fishes.

## Results

### Muscle growth dynamics vary greatly between closely related danionins

To assess lifelong muscle growth capacities in the danionins and killifish, we sampled their skeletal muscle at various critical developmental stages, including the early larval (pSB), late larval (DR), early juvenile (J), early adult (A) and mid adult stages (A+) (Fig. 1A&B, S1A)^30^. Muscle cross sections were stained for myofibers and ECM (Fig. S1B), then imaged and analyzed using a custom, semi-automated muscle morphometry analysis workflow to segment and morphometrically assess each myofiber in the trunk musculature (Fig. 1C-F, S2C). The analyses revealed that while all species exhibit marked increases in both myofiber count and myofiber size during postembryonic development, divergence in hyperplastic growth trajectories is the primary contributor to differences in growth capacities between species (Fig. 1E), with *Danionella* exhibiting the lowest hyperplastic growth capacity, giant danio exhibiting maximal hyperplastic muscle growth, while zebrafish and killifish exhibit moderate hyperplastic muscle growth capacities. Importantly, while hyperplastic muscle growth in zebrafish and *Danionella* primarily occurs before early juvenile phase, giant danio and killifish continue to undergo significant hyperplastic muscle growth as juveniles (193% and 61.2% increases in myofiber count respectively between J and A), only approaching a plateau at the early adult phase. This increase in hyperplastic growth capacity during juvenile giant danio and killifish development is further associated with their ability to increase total muscle mass post-maturity (Fig. S2A), which is lacking in zebrafish and *Danionella*, highlighting the importance of hyperplasia in supporting continuous muscle growth in teleosts.

**Fig 1.**
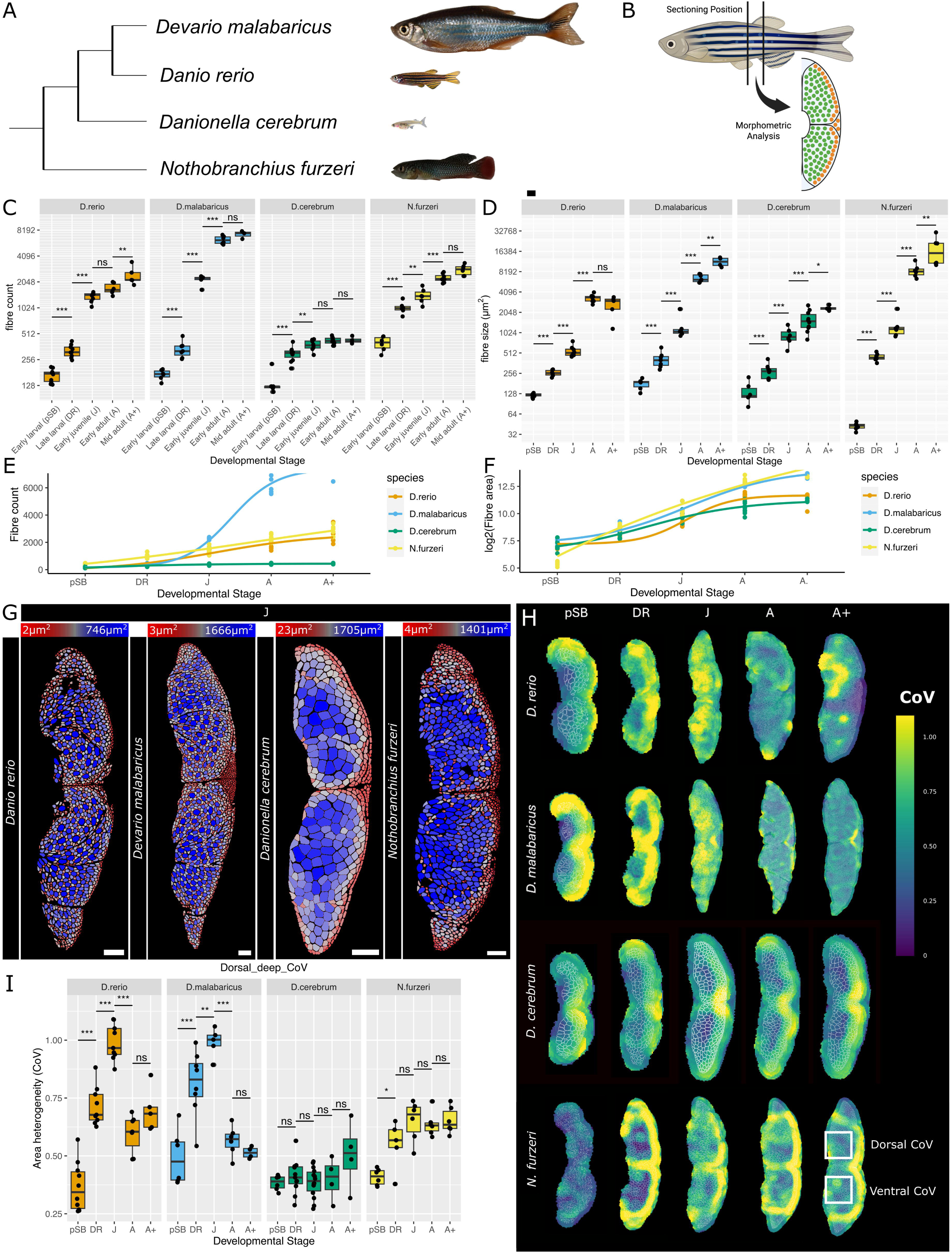
Alterations in mosaic hyperplasia contribute to major differences in muscle growth capacities between teleosts. (A) Danionins have drastic differences in body size, as shown by representative images of early adults in each examined species. Scalebar: 4 mm. Phylogenies were adopted from^79–81^. (B) Sections were obtained at a consistent position adjacent to the dorsal and anal fins, followed by muscle morphometry analysis. Green: fast myofibers, orange: slow myofibers. (C) Quantification of myofiber count in either the left or right half of the trunk muscle in each transverse section. (D) Mean area of the largest 100 myofibers in each transverse section. Note that statistical analysis was performed using the area of all 100 myofibers per sample. (E) Level of hyperplastic muscle growth between each stage, represented as log2 fold-change, fitted with sigmoidal curves. (F) Level of hypertrophic muscle growth between each stage, represented as log2 fold-change, fitted with sigmoidal curves. (G) Representative images of fibre area distribution heatmaps in each species at early juvenile stage, when mosaic hyperplasia maximizes. Smallest myofiber in each image is pseudo-colored in red and largest colored in blue. Scalebar: 100 µm. (H) Representative images of area variation heatmaps: regions undergoing hyperplasia has large variation between small, newly synthesized fibers and larger, older fibers, resulting in large area variation, colored by yellow. At larval stages, highest variation occurs at myotome boundaries, suggesting stratified hyperplasia as the main driver of growth. However, in zebrafish and giant danio at the early juvenile stage, highest variation occurs deep inside the myotome, suggesting mosaic hyperplasia as the key driver of growth. (I) Quantification of CoV deep inside the dorsal myotome. Killifish has reduced mosaic hyperplasia during development, whereas *danionella* portrays complete absence of mosaic hyperplasia. Stats: linear models (for C&I) and linear mixed models (for D), followed by pairwise comparisons using estimated marginal means with Sidak-adjusted p-values. ns > 0.05, * < 0.05, ** < 0.01, *** < 0.001.

### Alterations in hyperplastic growth patterning contribute to species-specific growth differences

Spatial patterning of hyperplastic muscle growth has previously been implicated to affect muscle growth capacities in teleosts^31–34^, though this effect was never quantified and compared across species. To visualize where small, newly synthesized myofibers localize during teleost development, we pseudo-colored each myofiber by area, with the smallest myofibers in each sample shown in red (Fig. S1C). The resulting heatmaps reveal that small myofibers initially emerge solely on the edges of myotomes in early larval teleosts, a phenomenon known as stratified hyperplasia. However, in all species except the *Danionella*, small myofibers also emerge deep in the myotome starting from late larval stage. This myofiber synthesis pattern, known as mosaic hyperplasia, then maximizes at the early juvenile stage in zebrafish, giant danio and killifish. Notably, *Danionella* did not exhibit small myofibers deep inside the myotome at any developmental stage, indicating a complete loss of mosaic hyperplasia, which likely underlies their reduced growth capacities (Fig. 1G).

While *Danionella* portrays a complete absence of mosaic hyperplasia, visual inspection of killifish myotome also suggested a reduction of small myofibers deep inside the myotome. To quantitatively assess the extent of mosaic hyperplasia, we developed a method to calculate the coefficient of variation (CoV) in myofiber area across the myotomes (Fig. 1H&I). By comparing CoV deep inside the dorsal and ventral trunk myotome across developmental stages and species, we found that zebrafish and giant danio both display high levels of mosaic hyperplasia during late larval to early juvenile stage, whereas killifish has greatly reduced mosaic hyperplasia (Fig. 1I, S2B). Additionally, in line with earlier observations, CoV in *Danionella* never increases deep inside the myotome during development, supporting complete absence of mosaic hyperplasia. Taken together, these observations indicate that mosaic hyperplasia is a major driver of muscle growth in teleosts, contributing to the divergence of growth capacities between species.

### MuSC-specific expression of ECM genes inhibits activation and hyperplastic muscle growth

To examine the molecular mechanisms underlying the attenuation of hyperplastic muscle growth in juvenile zebrafish, we next performed a scRNA-seq experiment using cells isolated from trunk muscle tissue of early juvenile and early adult zebrafish. After normalization, dimensionality reduction and unsupervised clustering, we manually annotated each cluster by referencing a list of cell marker genes, in addition to performing gene set enrichment analysis (GSEA) using cluster markers to identify enriched cell types (Fig. S3A-D). We subsetted the dataset for muscle stem and progenitor cells only, then re-clustered these myogenic cells to find 4 unique clusters representing quiescent MuSCs (highly expressing quiescent markers *pax7a/pax7b*), activated MuSCs (lower *pax7a/pax7b* expression and higher ribosomal gene expression), committed myoblasts (upregulating *myog*) and myocytes (upregulating *actc1b*), respectively (Fig. 2A&B), which differentiate in this order according to trajectory analysis (Fig. 2C) and in line with known myogenic differentiation processes^21,35^. Differential expression analysis (DEA) between myogenic cells in juvenile and adult zebrafish followed by GSEA revealed genes associated with ECM receptor interactions to be upregulated in adult myogenic cells (Fig. 2D&E), suggesting their potential function in mediating the cessation of hyperplastic muscle growth in adult zebrafish.

**Fig 2.**
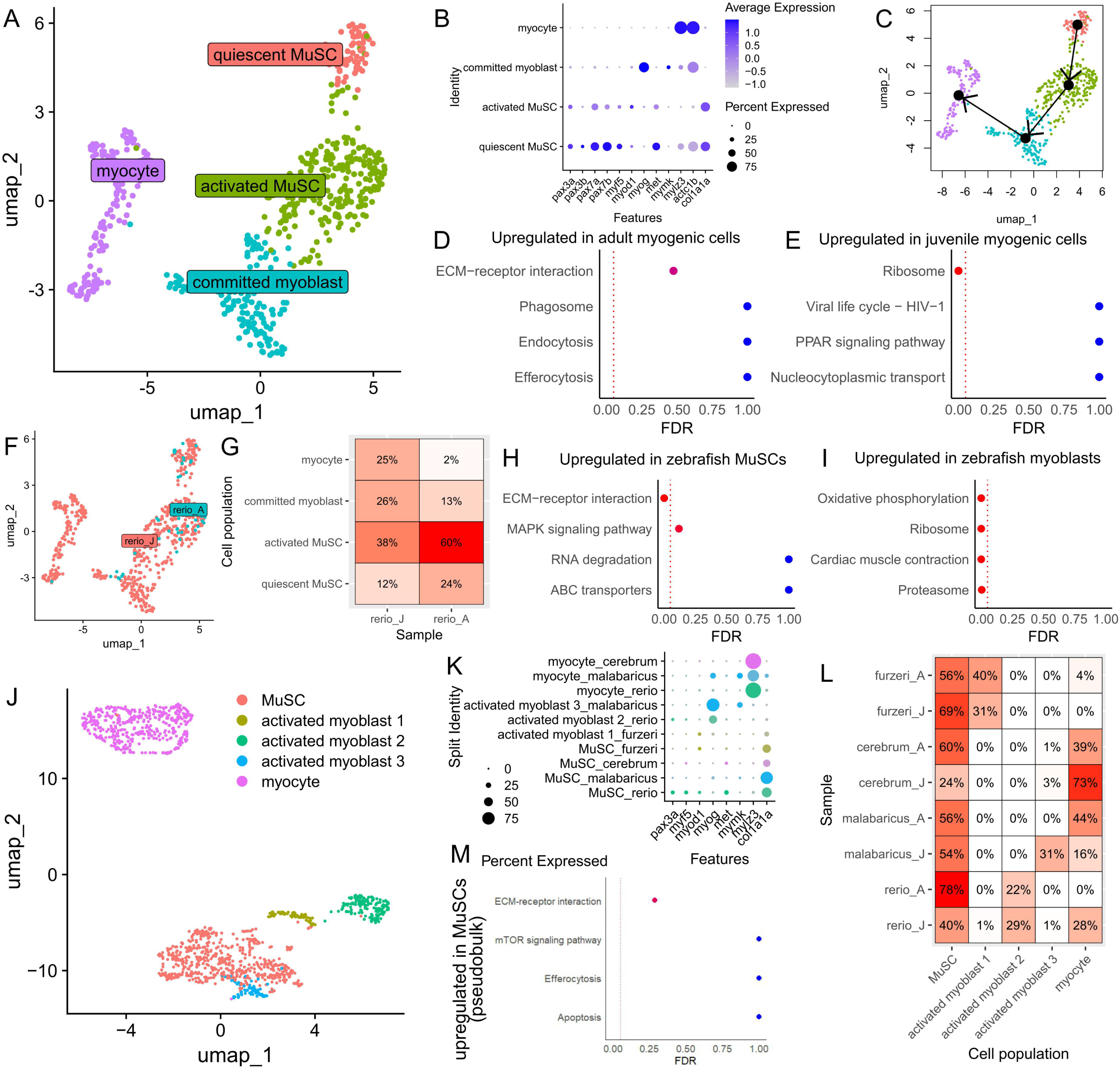
scRNA-seq reveals inhibitory effect of MuSC-expressed ECM genes on cell activation and hyperplastic muscle growth. (A) UMAP of myogenic cell clusters. After subsetting the full dataset, four myogenic clusters were found. (B) Dot plot showing expression of key markers in each cluster. Notably, quiescent MuSCs are marked by *pax3a*, *pax7a*, *pax7b* and *met* expression, activated MuSCs are marked by lower *pax7* expression, committed myoblasts are marked by *myog* expression and myocyte are marked by *actc1b* and *mylz3* expression. (C) Trajectory analysis using Slingshot revealed a differentiation trajectory starting from quiescent MuSCs, followed by activated MuSCs and committed myoblasts, ending with myocytes. (D&E) GSEA reveals upregulation of ECM-receptor interactions pathway and downregulation of ribosomes in adult myogenic cells compared to juvenile myogenic cells. (F) UMAP demonstrating the contribution of each sample to myogenic cells. (G) Heatmap portraying the percentage of each myogenic cell type that each sample contributes to. (H&I) GSEA reveals upregulation of ECM receptor interactions in MuSCs compared to myoblasts, suggesting their role in promoting quiescence. In myoblasts, oxidative phosphorylation, ribosomes, muscle contractile apparatus (annotated as cardiac muscle contraction in KEGG) and proteasomes are upregulated. (J) UMAP portraying integrated datasets for all 4 species combined, subsetted for myogenic populations. (K) Expression of key marker genes in each cell population in integrated dataset. (L) Heatmap showing proportion of each myogenic cell type in each sample. Note that each species has its own characteristic myoblast population. (M) Pseudo-bulk analysis supports an evolutionarily conserved upregulation of ECM-receptor interactions pathway in MuSCs compared to myoblasts and myocytes.

To better understand the cause of ECM gene upregulation, we calculated the proportion of each myogenic cell type in each sample (Fig. 2F&G), which revealed that adult zebrafish exhibits increased number of quiescent or activated MuSCs and fewer myoblasts/myocytes, in line with their reduced hyperplasia capacity. Importantly, DEA followed by GSEA between MuSCs and myoblasts showed downregulation of ECM-receptor interactions associated genes (*col1a1b, col1a2, col4a1, col4a2, col6a1, lamb1a, lamc1, hspg2* and *dag1*) during MuSC activation and differentiation, while oxidative phosphorylation, translational machinery (ribosomes) and muscle contractile proteins are upregulated (Fig. 2H&I). These results suggest that MuSC-specific expression of ECM genes in zebrafish have inhibitory effects on MuSC activation, therefore reducing myoblast/myocyte number and suppressing hyperplasia.

To further examine if the ECM plays evolutionarily conserved growth-regulatory roles in other teleost species, we performed additional scRNA-seq experiments to study the muscles of early juvenile and early adult *Danionella*, giant danio, and killifish, respectively. Next, using reciprocal best-hit BLAST to identify orthologs between species^36^, we integrated the dataset from all four species into a single dataset. We transferred cluster annotations from the earlier zebrafish analysis into this integrated dataset, subsetted the sample for only myogenic cells, and reclustered these myogenic cells to yield a total of 5 myogenic cell clusters in the dataset (Fig. 2J&K). While the MuSC and myocyte populations from all four species cluster together, each species form their own characteristic myoblast population (Fig. 2J&L), with zebrafish, killifish, and giant danio each developing their own activated myoblast population, whereas *Danionella* does not have a myoblast population.

By performing DEA using all MuSCs against all myoblasts and myocytes in this integrated dataset, we again found upregulation of ECM-receptor interactions in the MuSCs, suggesting MuSC-specific expression of ECM genes to be a conserved phenomenon in all examined teleosts. We also tested this finding by pseudo-bulking cells by species and cell cluster, followed by DEA and GSEA, which again revealed ECM-receptor interaction-associated genes (*col6a1, col6a2, itga5, itga9, thbs2a, sdc4* and *lamc1*) to be upregulated in MuSCs (Fig. 2M), supporting their roles in defining MuSC identity in teleosts. Note that due to challenges in identifying 1:1 orthologs for collagen genes across species, these collagen genes are greatly underrepresented in this new list.

Since our scRNA-Seq analysis repeatedly identified upregulation of collagen and thrombospondin genes in MuSCs, which have previously been linked to MuSC activity and myopathies^37–43^, we followed up on these results by performed in situ HCR to label *pax7a^+^*, *col1a1a^+^*, and *thbs2a^+^*cells in juvenile teleost muscle sections. Additionally, we co-labelled proliferative cells using phospho-histone H3 antibodies (pH3). We used a custom, semi-automated workflow to segment and identify each MuSC by their DAPI label and *pax7a* expression, then assessed the correlation between ECM gene expression and proliferative status by comparing their *col1a1a* expression, *thbs2a* expression, and pH3 staining levels (Fig. 3A, S3H). To simplify comparisons, we categorized *col1a1a* and *thbs2a* expression levels into 3 equally sized bins according to their intensities – low, medium, and high. We then statistically assessed the correlation between *col1a1a/thbs2a* and pH3 intensities using linear mixed models, taking each fish as the random effect. These models revealed significant association between ECM gene expression and proliferative status in MuSCs (Fig. 3B). Following this, pairwise comparisons revealed a significant decrease in pH3 expression in both *col1a1a*-high and *thbs2a-*high MuSCs. Taken together, these results support the earlier finding that MuSC-specific expression of ECM genes inhibit their own activation, proliferation and ultimately hyperplastic muscle growth in teleosts.

**Fig 3.**
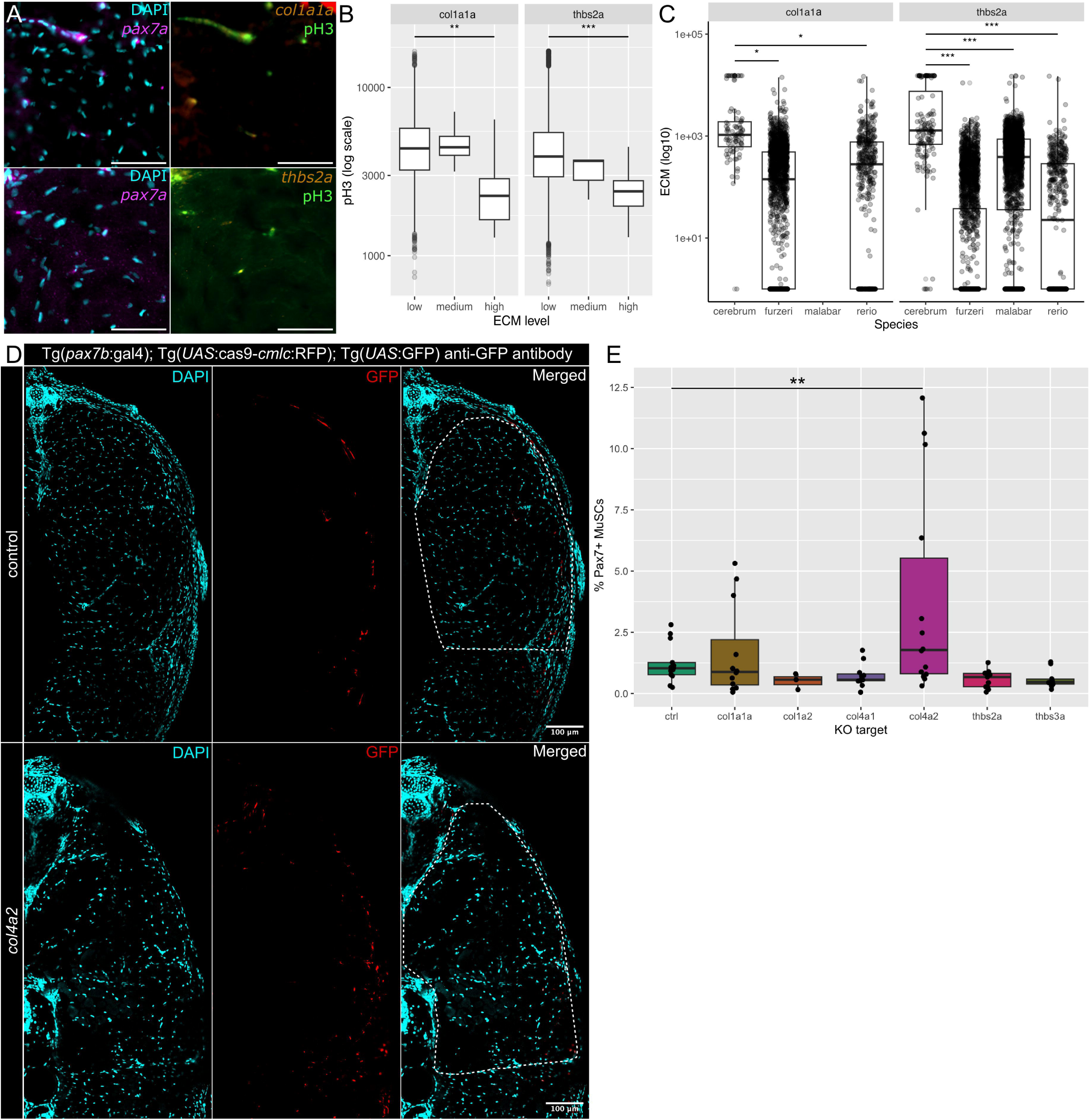
Expression and functional validation support an inhibitory effect of ECM genes on MuSC activity. (A) Representative images of in situ HCR on juvenile teleost muscle. The two images on each row each represent two channels of a four-channel image. The trunk musculature from an epaxial quadrant is examined in each sample. Scalebar: 50 µm. (B) *col1a1a* and *thbs2a* expression levels in DAPI^+^/*pax7a^+^* cells were categorized into 3 equally sized bins according to their intensities – low, medium, high, then compared against pH3 intensities. (C) *col1a1a* and *thbs2a* expression levels in DAPI^+^/*pax7a^+^* cells were binned by species and compared. (D) Representative images of stained muscle cross-sections from Tg(*pax7b*:gal4); Tg(*UAS*:Cas9-*cmlc*:RFP); Tg(*UAS*:GFP) fish injected with scrambled gRNA or *col4a2* gRNA. Cells are quantified from the epaxial quadrant of each sample, as outlined in the merged images. Scalebar: 100 µm. (E) Comparison of Pax7^+^ MuSC count, normalized against DAPI count, in each crispant. Stats: linear mixed models, followed by pairwise comparisons using estimated marginal means with Sidak-adjusted p-values. ns > 0.05, * < 0.05, ** < 0.01, *** < 0.001. Only significant pairwise comparisons are shown.

Next, to determine whether MuSC-specific ECM gene expression levels underly hyperplastic growth differences between species, we binned *col1a1a* and *thbs2a* expression levels in MuSCs by species. We then assessed differences in ECM gene expression using linear mixed models, once again taking individual fish as the random effect. While we could not design *col1a1a* HCR probes for giant danio, our results indicate that *Danionella* MuSCs significantly upregulate *col1a1a* compared to both zebrafish and killifish (Fig. 3C). Similarly, *thbs2a* expression is also significantly higher in *Danionella* MuSCs than the other three examined species. These observations align with the limited hyperplastic muscle growth capacity observed in *Danionella*, further supporting the role of ECM genes in inhibiting MuSC proliferation in teleosts.

Having shown the evolutionarily conserved association between collagen and thrombospondin gene expressions with MuSC activity in teleosts, we then turned to functionally validating their roles in MuSC regulation using a F0 zebrafish crispant screen. We targeted *col1a1a*, *col1a2*, *col4a1*, *col4a2, thbs2a*, and *thbs3a*, which represented some of the enriched ECM genes upregulated in inactive MuSCs. We injected gRNAs targeting each of these genes in turn into Tg(*pax7b*:gal4); Tg(*UAS*:Cas9-*cmlc*:RFP); Tg(*UAS*:GFP) zebrafish at the one-cell stage to achieve MuSC-specific deletion. Muscle cross-sections from crispants were examined at early juvenile stage to determine Pax7^+^ MuSC count using an automated cell counting workflow (Fig. 3D). We then used a linear mixed model to compare the normalized MuSC count in each crispant against control fish injected with scrambled gRNA, taking individual fish as the random effect. Pairwise comparison revealed a significant increase in Pax7^+^ MuSC prevalence in the *col4a2* crispants (Fig. 3E), supporting its role in regulating MuSC dynamics.

## Discussion

A number of teleost fishes with available genome sequences serve as vertebrate models for comparative biology and translational research^44,45^, with zebrafish being commonly used to study skeletal muscle development, growth, and repair^23,24^. In this study, we demonstrated that the spatial patterning of myofiber synthesis during hyperplastic muscle growth is a major contributor to the divergence of growth capacities between teleosts. Even among the closely related danionins, mosaic hyperplasia capacities during larval and juvenile development could range from near-zero in *Danionella*, to extensive in the zebrafish and giant danio, suggesting a highly adaptable process. At the molecular level, hyperplastic muscle growth is driven by MuSC activation, proliferation, and differentiation, which is further regulated by their autonomous expressing of ECM genes, a mechanism we found to be conserved across the examined teleosts. Importantly, by MuSC-specific depletion of the *col4a2* gene, we altered MuSC dynamics in zebrafish, suggesting the ECM pathway to be a plausible target for mediating muscle growth in vertebrates.

In line with our findings, past studies have implicated various ECM genes in mediating MuSC activity. For example, culturing mammalian MuSCs with collagen V was shown to repress their differentiation, whereas depletion of a critical subunit of collagen V leads to the rapid loss of the MuSC pool^37^. Similarly, *Col6a1* deletion in mice impairs MuSC self-renewal and muscle regeneration^46^, supporting another study that demonstrated collagen VI to exhibit moderate, inhibitory effect on MuSC activation in mammalian myogenic cell lines^38^. Furthermore, a study in larval zebrafish had described *col1a2* expression in muscle stem and progenitor cells, tenocytes, and xanthophores^47^. When these *col1a2-*expressing cells were ablated using the metronidazole/nitroreductase system, the authors observed compromised muscle regeneration in larvae, further supporting a critical role of collagens in maintaining the MuSC pool. A recent study in trout found simultaneous upregulation of *col5a3a* and downregulation of *col6a3* in muscle stem and progenitor cells during development, as well as differential expression of numerous other ECM genes, potentially suggesting antagonistic roles of different ECM genes in mediating MuSC dynamics^48^. While we did not observe a similar phenomenon, further characterization of the precise effect of each ECM gene on MuSC dynamics could be an important future direction.

Historically, mosaic hyperplasia has been hypothesized to drive indeterminate growth in large teleosts, whereas smaller teleosts such as zebrafish and guppies were believed to undergo stratified hyperplasia only^49,50^. However, this misconception likely arose from inadequate sampling of late larval and early juvenile teleosts, when mosaic hyperplasia is most prominent. Here, we demonstrate that although zebrafish is a small, determinate grower, its mosaic hyperplasia capacity at early juvenile stage is equivalent to the giant danio that’s believed to represent larger, indeterminate growers. Thus, mosaic hyperplasia is not necessary for indeterminate growth^34,51^. Despite falsifying the idea that mosaic hyperplasia drives indeterminate muscle growth in teleosts, this study has demonstrated that the relative contribution of stratified and mosaic hyperplasia can be modulated, with killifish portraying reduced mosaic hyperplasia (despite having greater body size than zebrafish) and *Danionella* portraying no mosaic hyperplasia. Thus, mosaic hyperplasia is an important and highly modifiable contributor to the divergence of muscle growth capacities between teleost species.

Beyond the spatial patterning of myofiber synthesis, growth capacity in teleost is also influenced by alterations in developmental timing (heterochrony), a phenomenon that we have not investigated in detail. Notably, while all three danionin species undergo a phylotypic larval phase with comparable growth duration (∼1 month), their juvenile developmental durations differ markedly. *Danionella*, which has lowest muscle growth capacity, also undergoes the shortest juvenile developmental duration of ∼ 1 month before reaching sexual maturity. In contrast, zebrafish and giant danio undergo ∼2 months and ∼11 months of juvenile development respectively. The difference in developmental duration correlates with muscle growth capacities in each danionin species. Indeed, although zebrafish and giant danio have similar spatial patterning of hyperplasia at any single developmental stage, the prolonged developmental duration in giant danio ultimately yields increased total muscle growth, resulting in larger body size. Thus, heterochrony is also an important and effective evolutionary strategy for mediating divergence in muscle and body sizes between species.

In our current study, we functionally validated the effect MuSC-specific expression of *col4a2* on MuSC dynamics. However, we did not observe similar phenotype when disrupting the other ECM genes. There are multiple challenges that may have resulted in these negative results. Firstly, ECM gene expression in MuSCs are self-regulatory via a negative feedback loop to maintain ECM stiffness and cell function^52^. Under certain pathological conditions, such negative feedback could be disrupted^53–56^, though this contributes to additional complexity to the functional validation. Secondly, a truly MuSC-specific modulation of gene expression is difficult to achieve, as critical MuSC markers such as *pax3a*, *pax3b*, *pax7a* and *pax7b* are also expressed in brain, spinal cord and pigment cells, which could exert additional effects on MuSC dynamics. Thirdly, fibroblasts are also critical contributors of ECM proteins and potentially help establish the ECM niche that regulates MuSC dynamics^57^, which will need to be better understood to design effective strategies to modulate ECM dynamics. In future studies, overcoming these challenges will be important to enabling the modulation of muscle growth dynamics in vertebrates by targeting the ECM pathway.

## Online Methods

### Fish husbandry

All animal works were approved by the Monash Animal Ethics committee. Zebrafish of the TU strain were raised according to standard husbandry procedures. Shortly, juvenile and adult zebrafish were raised in circulating aquarium system at 26.5°C, with a 14h light – 10h dark cycle. Eggs were kept in E3 embryo medium (5 mM NaCl, 0.17 mM KCl, 0.33 mM CaCl, 0.33 mM MgSO4, 0.00004% [v/v] methylene blue in water, pH 7.2) in petri dishes until up to 7 dpf, after which they are fed with paramecium twice daily until they reach up to 10 dpf. Larvae aged between 10-30 dpf were fed with *Artemia salina* once daily in addition to the paramecia. Juvenile and adults above 30 dpf were fed with *Artemia salina* and appropriately sized pellets once daily. Zebrafish was housed in 4L tanks, to a maximum concentration of 5 fish per litre water. The following transgenic lines were previously generated and used here: (1) TgBAC(*pax3a*:GFP); TgBAC(*meox1*:mCherry) (unpublished), (2) Tg(*pax7b*:gal4); Tg(*UAS*:Cas9-*cmlc*:RFP); Tg(*UAS*:GFP)^58^. Additionally, we generated MuSC-specific knockouts for *col1a1a*, *col1a2*, *col4a1*, *col4a2, thbs2a* or *thbs3a*. Husbandry of *Danionella cerebrum* was performed according to published protocols as well^59,60^.

Giant danio (*Devario malabaricus*) was obtained from Aquarium Industries Pty Ltd. (Epping, Victoria) and housed on aquarium systems with identical parameters to zebrafish. However, the following modifications were made to account for their larger sizes: juvenile and adult fish larger than 4 cm SL were housed in custom-made 160L tanks equipped with wavemakers, at a maximum density of 4 litre water per fish. Additionally, giant danio greater than 4 cm SL also received live blackworms once a day to supplement nutritional needs. Green, sturdy, plastic plants placed in the 160L tanks to stimulate breeding, the laid eggs are then collected from the plastic plants and kept in E3 medium in petri dishes until up to 5 dpf before they are fed.

African Killifish (*Nothobranchius furzeri*) strain MZCS_08/122 was housed according to published protocols^61^, with the following alterations: Glass beads were used in place of sand. Freshly collected killifish eggs were bleached by incubating in 5% sodium hypochlorite solution for 2 min, followed by sterilised system water for 2 min, followed by 5% sodium hypochlorite solution for 2 min, followed by additional 2 washes in sterilised system water, each for 2 min, then stored in killifish embryo medium (0.2% methylene blue, 10 mg/L gentamicin in system water) until black eyes are clearly visible. These embryos are then transferred onto moist filter paper containing killifish embryo medium instead of coconut fiber, until their eyes turn completely golden, signifying the ready-to-hatch stage. These embryos are then subjected to hatching in 4°C sterilised system water with a continuous supply of airflow. Fish under one week post hatch are fed with paramecia and artemia, following which they are fed with artemia, fish pellets, and live blackworms.

We relied on anatomical features in developing teleosts to accurate identify developmental stages in examined teleost, in accordance to the zebrafish normal table of development^30^. Four developmental stages from each species were characterized, including early larval stage (pSB) characterized by swim bladder inflation, late larval stage (DR) characterized by dorsal fin ray formation, early juvenile stage (J), characterized by scale formation and early adult stage (A) characterized by sexual maturity. An additional mid-adult stage was examined, taken at a midpoint between maturity age and expected lifespan in each species respectively. This corresponds to 1 year post fertilization (1 ypf) in zebrafish, 2 ypf in giant danio, 22 weeks post hatch (22 wph) in killifish and 22 wpf in *Danionella*. Larval and adult fish were anaesthetized in E3 embryo medium (or system water) containing 0.02% Tricaine Methane sulfonate (MS-222) until movement ceases. For Euthanasia, tricaine concentration was increased to 0.4%. A random mix or males and females was included for all species and developmental stages.

### Immunohistochemistry

Muscle blocks under 10 mm thickness and adjacent to the dorsal fin were dissected and fixed overnight in 4% PFA in PBS at 4°C, then washed into PBST. For vibratome sectioning, muscle blocks were then embedded in 4% molecular grade agarose (Meridian biosciences, BIO41025), then sectioned using Leica VT1200s Vibratome to obtain 200 μm transverse sections. Vibratome sections were stored in PBST at 4°C short-term or in 100% Methanol at -20°C long-term. For cryosectioning, fixed muscle blocks were washed serially into 10%, 20% and 30% sucrose in PBST at 4°C. Tissue blocks were subsequently embedded in Optimum Cutting Temperature Compound (OCT, Tissue-Tek), then snap-frozen in 2-methybutane chilled with liquid nitrogen. Cryoembedded samples were stored at -80°C until sectioning. 12 μm cryosections were obtained using Leica CM3050 S cryostat, captured on SuperFrost™ Plus slides (epredia), then stored at -80°C.

Next, all cryosections and vibratome sections were incubated in goat serum blocking solution (2% goat serum, 1% DMSO, 1%BSA, 0.1% Tween 20 in PBS), followed by 4°C overnight incubation in primary antibodies diluted in appropriate blocking solutions. These primary antibodies include anti-GFP (1:300, Invitrogen, A11122) and anti-pH3 (1:300, Sigma, 06-570). Next, sections were washed into PBST, then incubated in secondary antibodies and dyes diluted in corresponding blocking solutions for 4h (cryosections) or overnight at 4°C (for vibratome sections) respectively, then washed into PBST. The following dyes and secondary antibodies were used: DAPI (1:300), Phalloidin 555 (1:400, Invitrogen, A30106), Phalloidin 647 (1:400, Invitrogen, A22287), WGA-fluorescein (1:300, VectorLabs, FL-1021-5), WGA-rhodamine (1:300, VectorLabs, RL-1022-5) and various Alexa Fluor antibodies (Invitrogen) targeting corresponding primary antibodies.

### Microscopy

For imaging, all vibratome sections were mounted in 1% low melt agarose plates and imaged via Leica SP8 multiphoton (confocal) microscope controlled by Las X software. Cryosections on slides were mounted in 80% glycerol and coverslips, then imaged with Leica thunder deconvolution microscope (widefield) controlled by Las X software, including image deconvolution. All images were captured at a resolution of 512 x 512 pixels or 1024 x 1024 pixels per tile and bit depths of 8-16 bit. Confocal images were captured with 2-4 line averaging or line accumulation, with 1 airy unit pinhole size.

### Image analysis

Muscle morphometry analysis was performed using custom written scripts and macros in FIJI. Briefly, this involved an initial image segmentation using ilastik^62^, Labkit^63^ or Cellpose^64^ to identify myofibers, followed by manual correction to accurately obtain the full list of myofibers. Morphometric parameters of these myofibers were then quantified and visualized. Furthermore, variations (CoV) in area between neighboring myofibers were calculated and visualized using custom written Julia scripts. The CoV was computed at each point by considering all cells whose center was within a window radius *r*. The window radius which was taken to be three times the mean Feret diameter of each cell in the image. For a given point, the CoV was calculated as the mean divided by standard deviation of cell areas within the window radius.

Cell counting was performed in FIJI2 (ImageJ2) using custom written macros. Briefly, images were first manually cropped to remove signals outside of the trunk myotome. Next, all relevant channels were segmented using Labkit^63^ to identify DAPI^+^ cells and assess expression of stem cell or proliferation markers. For in situ HCR, we used control slides stained with only secondary antibodies to assess background intensities, which were used to correct for the expression levels of ECM genes and pH3. The number of cells that express different markers were then collated for statistical analysis. All codes can be made available upon request.

### RNA extraction & bulk RNA-sequencing

To generate a giant danio reference transcriptome, RNA was isolated from freshly dissected muscle tissue. Specifically, we extracted RNA from early larval (pSB), late larval (DR), and early juvenile (J) tissues using RNeasy Kits for RNA Purification (Qiagen) according to manufacturer’s protocol, or using TRI reagent (Invitrogen, AM9738) as follows. The tissue was placed in 1 ml TRI reagent and homogenized approximately 10 times using a Dounce homogenizer. The homogenized sample was incubated at room temperature for 5 minutes to ensure complete dissociation of nucleoprotein complexes. Then, 200 μl of chloroform was added to the sample, followed by a 15-second vortex, and the sample was incubated at room temperature for 3 minutes. The sample was then centrifuged at 12,000 g for 20 minutes at 4°C, and the top aqueous phase containing RNA was collected. This was mixed with an equal volume of isopropanol and incubated at room temperature for 10 minutes. After centrifugation at 13,000 g for 20 minutes at 4°C, the supernatant was removed, and the pellet was resuspended in 1 ml of 75% ethanol. The RNA was then pelleted again by centrifugation at 13,000 g for 10 minutes at 4°C. The supernatant was discarded, and the pellet was air-dried for 10 minutes before being resuspended in nuclease-free water (Qiagen, cat #129117) and stored at -80°C.

Quality control (QC) was conducted by Micromon Genomics (Monash University, Clayton) to ensure an RNA integrity score greater than 8.0. Once the QC was passed, RNA samples were sent to the Beijing Genomics Institute (BGI) for sequencing using DNBSEQ Eukaryotic Strand-Specific Transcriptome Resequencing technology (short-read sequencing) and PacBio HiFi sequencing technology (long-read sequencing).

### Giant danio reference transcriptome assembly

Short- and long-read bulk RNA-seq data were processed using command-line interface (CLI) tools on the M3 (MASSIVE) server and custom R scripts on local systems. For short-read data, quality control was performed using FastQC (https://www.bioinformatics.babraham.ac.uk/projects/fastqc/) and fastp^65^. For long-read data, we first combined the individual PacBio subreads in the raw subreads.bam file using CCS (https://ccs.how/), then processed them into clean reads using the BGI Full-Length RNA Analysis Pipeline (https://github.com/shizhuoxing/BGI-Full-Length-RNA-Analysis-Pipeline). Next, we assembled the short-reads using long-reads as a reference with Trinity^66^. To remove gene redundancies caused by transcript isoforms, we generated superTranscripts^67^ from the Trinity output. The completeness of the reference transcriptome was then validated using BUSCO^68^, revealing a completeness score of 94.6% (Single: 91.5%, Duplicated: 3.1%). Transdecoder (https://github.com/TransDecoder/TransDecoder) was used to identify all open reading frames and obtain protein sequences. BLASTn, BLASTx and BLASTp was then used to query giant danio transcripts against the zebrafish genome (Ensembl, https://www.ensembl.org/index.html) and proteome (UniProt, https://www.uniprot.org/). Results were compiled using custom R scripts, yielding an annotated giant danio transcriptome. Additionally, we manually modified the annotation of all giant danio mitochondrial genes by adding the “mt-” prefix, aligning with zebrafish genome annotations.

### Single-cell RNA sequencing (scRNA-seq)

Trunk muscle tissues, containing both slow and fast muscles, were dissected at a region between the dorsal and anal fins of juvenile and adult fish. These tissues were then freshly dissected and minced using scalpel blades, then incubated in 2 ml of collagenase solution containing 1 mg/ml collagenase type II (Gibco™, cat #17101015) and 5% Fetal Bovine Serum (FBS, Gibco™, A5256701) in Hank’s Balanced Salt Solution (HBSS, Gibco™, cat #14025092) for 40-50 minutes at 32°C with shaking at 750 rpm. Samples were then centrifuged at 3500 rpm for 5 minutes at 4°C, after which the supernatant was removed. The pellets were resuspended in 2 ml of 0.25% Trypsin-EDTA (Gibco™, cat #25200072) and incubated at 30°C for 45 minutes with shaking at 750 rpm. After a second centrifugation at 3,500 rpm for 5 minutes at 4°C to remove the supernatant, the samples were resuspended in 2 ml of HBSS containing 5% FBS. The suspensions were filtered through 70 μm cell strainers (Falcon), centrifuged at 3,000 rpm for 5 minutes at 4°C to remove the supernatant, and resuspended in 2 ml of fresh HBSS with 5% FBS. The samples were filtered through 40 μm strainers (Falcon), centrifuged again at 3,000 rpm for 5 minutes at 4°C to remove the supernatant, and resuspended in 2 ml of fresh HBSS containing 5% FBS. Cell suspension was then processed via FACS to isolate live, single cells. 5 μl of the cell suspension was mixed with 5 μl of 0.4% Trypan blue (Invitrogen, T10282), loaded onto a Reichert Bright-line phase contrast hemocytometer, then examined under a Nikon Eclipse TS100 phase-contrast microscope to estimate the cell concentration. Average cell counts from five grids on the hemocytometer was used to estimate the cell density, and the suspension was adjusted to a concentration of ∼1000 cells/μl. The samples were kept on ice and submitted to Hudson Genomics (Hudson Institute of Medical Research, Clayton) or Research Institute for Microbial Diseases (RIMD) NGS core facility (The University of Osaka, Suita) for library preparation using GEM or GEM-X technology (10X Genomics), followed by sequencing with the NextSeq 2000 system according to protocol 1000000109376 v3 Nov2020 and processed with Bcl2fastq v2.20.2.422. ∼ 10,000 cells were sequenced per sample.

Following data acquisition, the raw reads were mapped using CellRanger V7.0.1 onto the GRCz11 zebrafish genome (ensembl), Nfu_20140520 killifish genome (ensembl), our custom giant danio transcriptome and *Danionella cerebrum* genome (unpublished), respectively. Downstream analysis was performed using the Seurat v5 workflow^69^. Quality control was performed independently for each species using nFeature_RNA and percent.mt as thresholds, followed by doublet removal using DoubletFinder^70^. To facilitate cross-species integration of scRNA-seq data, we utilized reciprocal best-hit BLASTp to identify one-to-one homologs between the 4 examined species^36^, using R and perl scripts provided by the authors. Custom R script was then used to fully integrate the dataset from each species. Data normalization was performed via SCTransform^71^. PCA was performed and elbow points were determined from scree plots, which were used to specify the number of dimensions to use for UMAP and clustering. Unsupervised clustering was performed at a resolution of 0.5. A list of marker genes was used to help annotate each cluster. Additionally, top 200 marker genes of each cluster were fed into FishEnrichr^72,73^ for gene set enrichment analysis (GSEA) to help infer cluster identities. Differential expression analysis was performed either performed using the FindMarkers function in Seurat (before pseudo-bulking) or DESeq2 package (after pseudo-bulking)^74^. Differentially expressed genes were queried using g:Profiler2 package in R^75^ to determine enriched gene sets and pathways. Trajectory analysis was performed using slingshot^76^. All code can be made available upon request.

### In situ HCR

In situ HCR was performed in line with the HCR™ RNA-FISH protocol for fresh frozen or fixed tissues sections provided by Molecular Instruments (Los Angeles, USA). Probes were designed according to published protocol^77^. Briefly, frozen tissue on slides were fixed with ice-cold 4% PFA for 15 min at 4°C, then washed through 50%, 70% and 100% ethanol, each for 5 min. Slides were washed again in 100% ethanol, immersed in PBS, digested with 10 ug/ml proteinase K for 5 min, followed by two quick PBS washes. Next, slides were equilibrated in hybridization buffer (Molecular Instruments) for 10 min in a 37°C humidified chamber, followed by overnight incubation in fresh hybridization buffer containing 4 nM of each probe set at 37°C. A coverslip was also added on top of each slide to prevent the slides from drying out. On the following day, coverslips were washed off, followed by a series of washes in probe wash buffers diluted in 5X SSCT (first 75% probe wash buffer, then 50%, 25% and 0%), each for 15 min at 37°C. Slides were washed in 5X SSCT again for 5 min, then incubated in amplification buffer (Molecular Instruments) for 30 min. To prepare hairpins, 6 pmol of hairpin h1 and h2 were independently heated to 95°C for 90 seconds, then cooled to room temperature in the dark. Hairpins were diluted in 100 ul of amplification buffer (Molecular Instruments) then added to each slide, followed by 48-hour incubation in a dark humidified chamber. Coverslips were placed on each sample again to prevent the slides from drying out. Lastly, coverslips were washed off using 5X SSCT, followed by three additional 5X SSCT washes on the slides, each for 30 min, then mounted using anti-fade mounting media and coverslips, followed by imaging. Probe set sequences can be found in Table S2.

### F0 CRISPR screen

We adopted a F0 CRISPR screen strategy to knock out candidate ECM genes including *col1a1a*, *col1a2*, *col4a1*, *col4a2*, *thbs2a* and *thbs3a* in zebrafish. In line with a published protocol^78^, three guide RNA targeting different exons of each gene were designed using IDT Predesigned Alt-R™ CRISPR-Cas9 guide RNA database (https://sg.idtdna.com/site/order/designtool/index/CRISPR_PREDESIGN), prioritizing those with high on-target scores and low off-target scores. All sequences are found in Table S1. Additionally, three scrambled crRNAs were included as controls (IDT cat #1072544, 1072545, 1072546). crRNA was heteroduplexed to universal tracrRNA in line with manufacturer’s protocol, then injected into freshly laid (<15 min old) eggs from Tg(*pax7b*:gal4); Tg(*UAS*:Cas9-*cmlc*:RFP); Tg(*UAS*:GFP) zebrafish to induce MuSC-specific deletion of these ECM genes.

## Acknowledgements

We thank all staff in the aquatic facilities at Monash University, NCVC and Stanford University for their assistance with teleost husbandry. We thank FlowCore for their assistance with FACS. We thank Hudson Genomics facility and RIMD NGS core facility for scRNA-Seq library preparation and sequencing. We thank Monash Micro Imaging (MMI) for its aid in microscopy, Monash Bioinformatics platform for its bioinformatic support and Monash eResearch for statistical support. We thank Dr Frank Tulenko for advice and assistance in histology. We thank Dr. Rebecca Dale for assistance in designing HCR probes. We thank Dr Wei Wang for providing the updated code for reciprocal best-hit BLAST. We thank Dr Braedan McCluskey for advice on danionin phylogeny. BioRender.com was used to create figures. Marco Podobnik thanks the Deutsche Forschungsgemeinschaft (DFG, German Research Foundation, project number: 549521547) for financial support. Michael Pan was supported by a postdoctoral research fellowship from the School of Mathematics and Statistics at the University of Melbourne. The Australian Regenerative Medicine Institute is supported by funds from the State Government of Victoria and the Australian Federal Government. The authors declare no conflicts of interest.

---

**Fig S1. Patterns of fiber synthesis during teleost development.** (A) Representative images of stages examined in each species. Mid-adult fish are not show as they appear similar to early adult fish. Scalebar: 4 mm. Phylogenies were adopted from^79–81^. (B) Representative images of muscle transverse sections from each species and stage. Muscle morphometry analysis in larval and juvenile fish was performed based on myofiber staining (phalloidin, green) on vibratome sections, whereas adult fish were analyzed based on ECM staining (WGA, red) on cryosections. (C) Representative images of myofiber area distribution heatmaps. Smallest fiber in each image is pseudo-colored in red and largest fiber colored in blue. Note that in zebrafish (*rerio*), giant danio (*malabaricus*) and killifish (*furzeri*), small, newly synthesized fibers first emerge deep inside the myotome at late larval phase, suggesting mosaic hyperplasia. In these three species, mosaic hyperplasia appears to peak at early juvenile stage, then declines by the early adult stage. Small, newly synthesized fibers are never observed deep inside the myotome of *Danionella cerebrum* (*cerebrum*), suggesting complete absence of mosaic hyperplasia. Scalebar: 100 µm.

**Fig S2. Muscle morphometry analysis.** (A) Quantification of total myofiber area in either the left or right half of the trunk muscle in each transverse section. Calculated by summing area of all fibers in each sample. (B) Quantification of CoV deep inside the ventral myotome is consistent with dorsal myotome, demonstrating reduced mosaic hyperplasia in killifish and no mosaic hyperplasia in *Danionella*. (C) Fiber area distribution in each stage. Note each graph contains fibers from all samples examined at that stage (for their respective species).

**Fig S3. scRNA-Seq and in situ HCR.** (A) UMAP portraying all cells sequenced from the zebrafish muscle. Note that muscle was dissected at a position adjacent to the dorsal and anal fins, free from any internal organs. (B) Dot plot showing expression of key markers in each cluster. (C) UMAP demonstrating the contribution of each sample to the different cell types. (D) Heatmap showing proportion of each cell type in each sample. (E) Integrated datasets containing cells from all four species. (F) Contribution of each sample to each cell type, UMAPs are separated by species. (G) contribution of each sample to the myogenic cell subset. (H) Representative images of in situ HCR from juvenile zebrafish. Scale bar: 100 µm.

