## Supplementary figures and images for "Divergence in skeletal muscle growth by differential spatial hyperplastic patterning in teleost fishes"

### Fig. S1

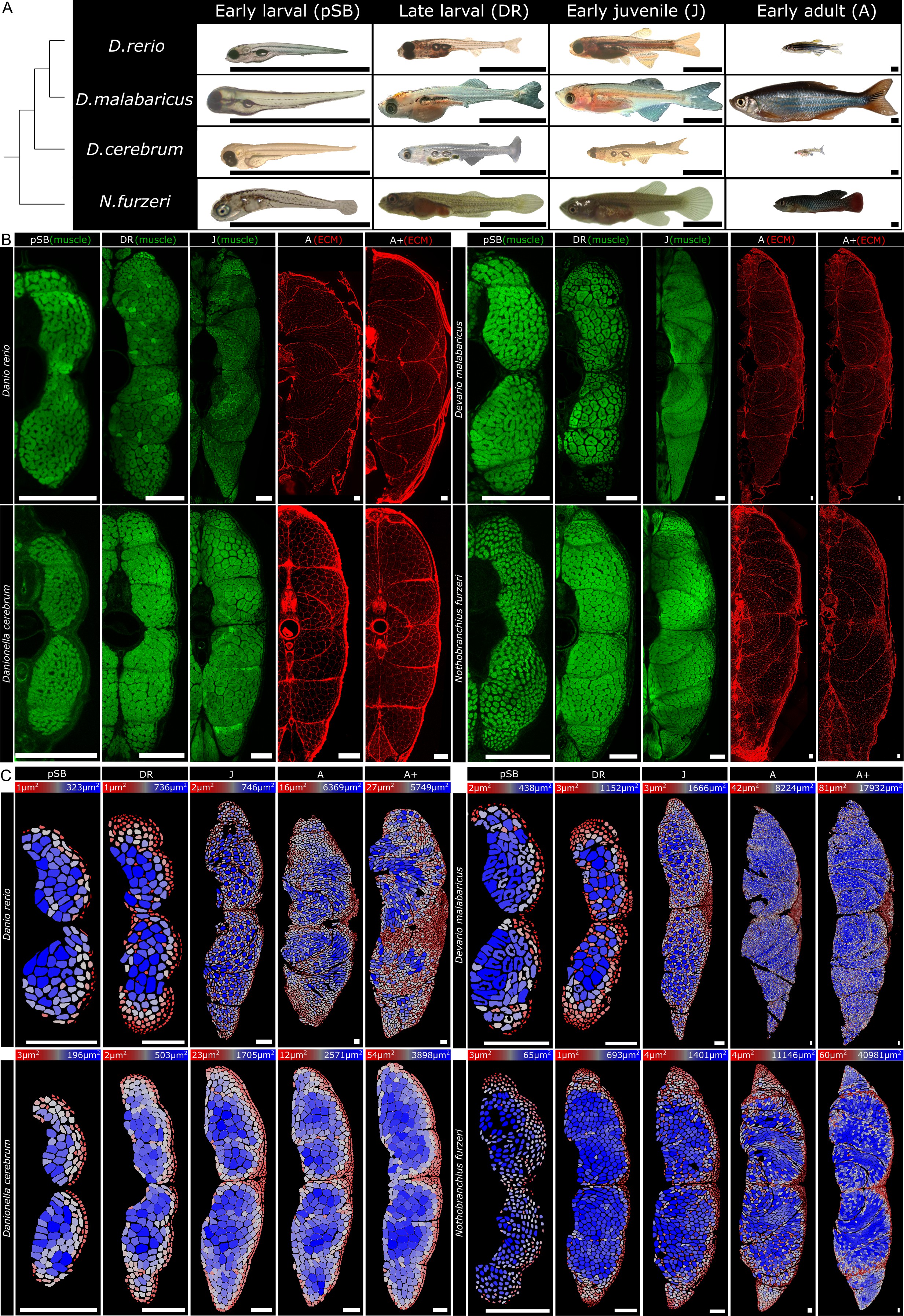

### Fig. S2

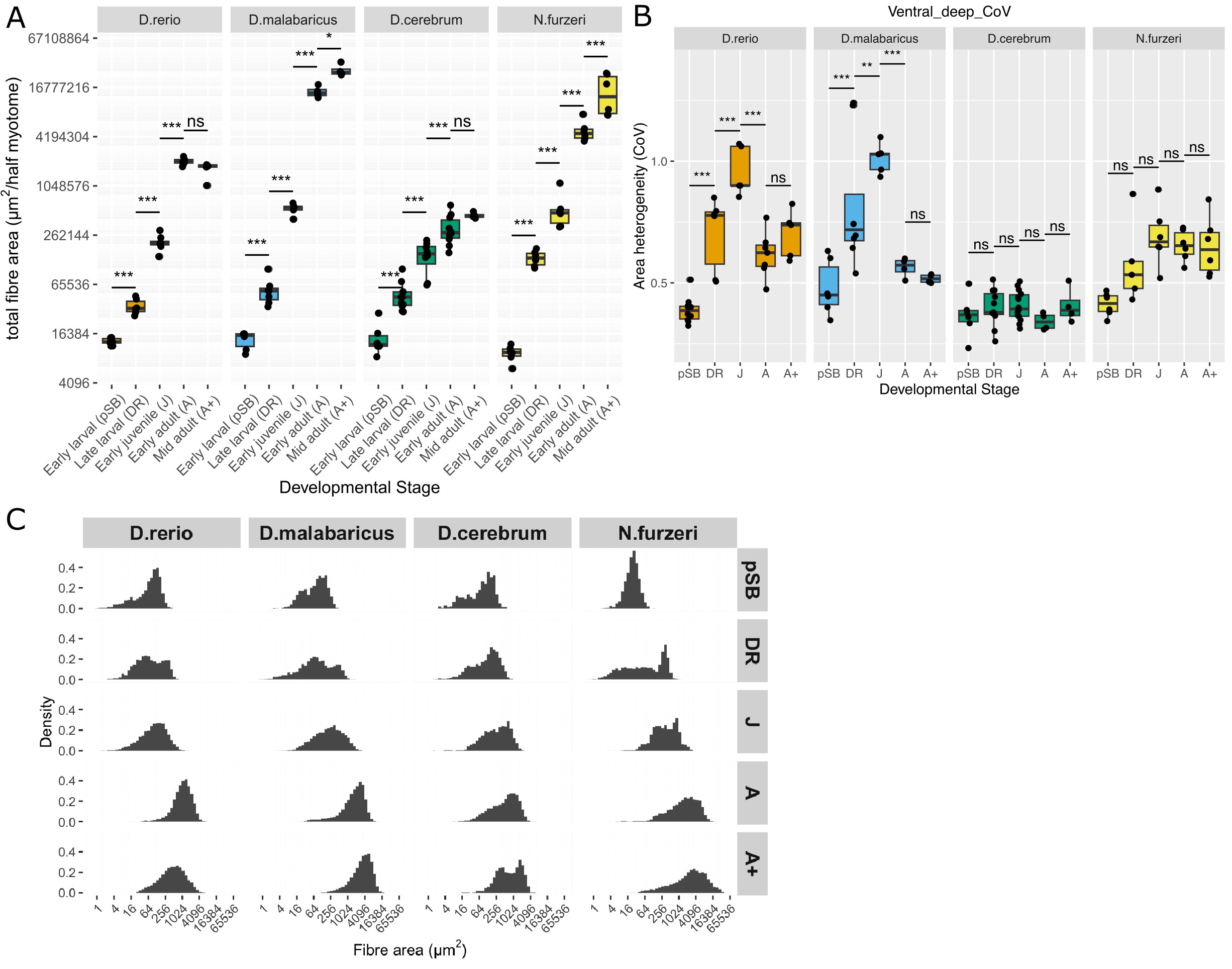

### Fig. S3

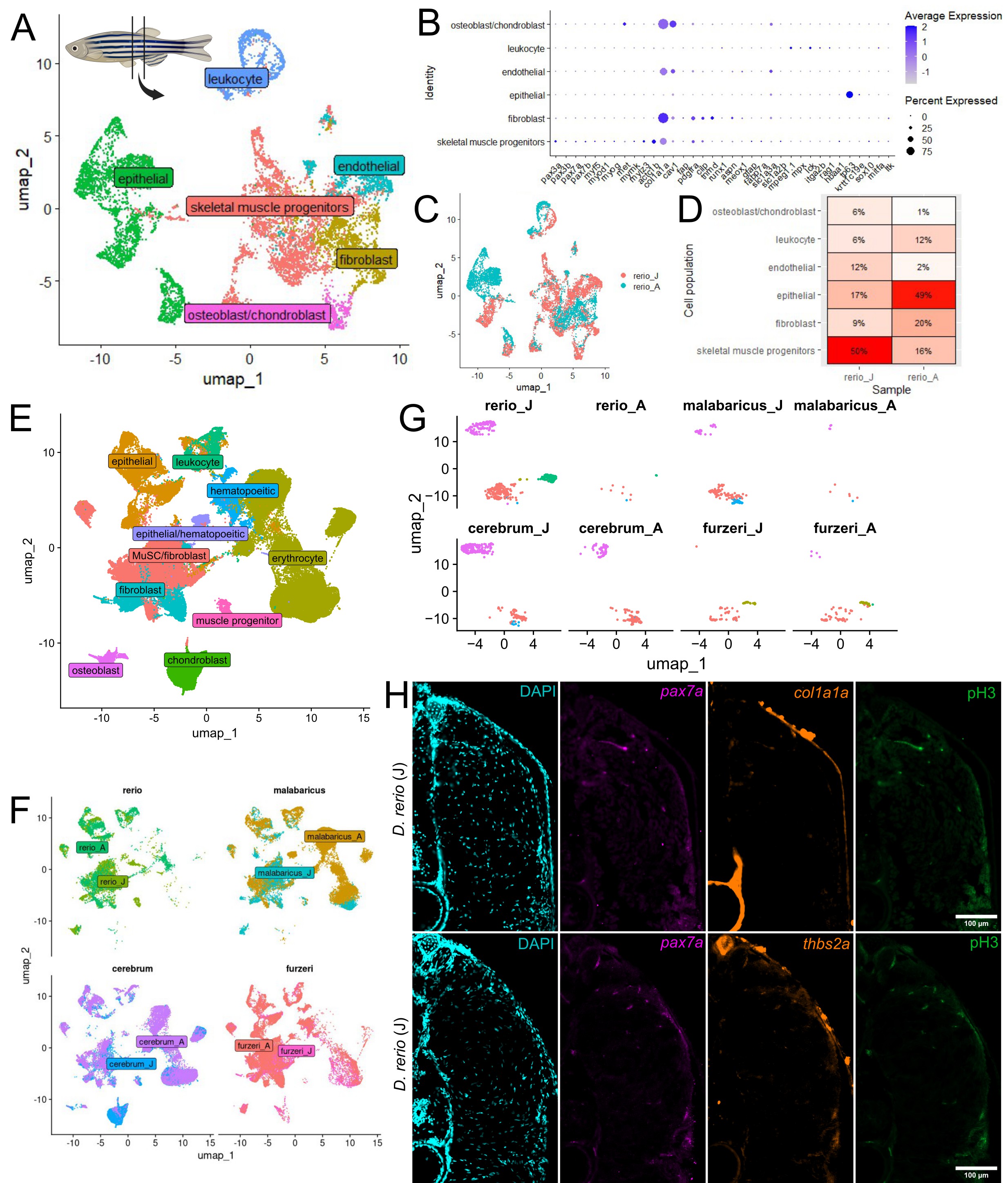
